# Additive and synergistic effects of arbuscular mycorrhizal fungi, insect pollination and nutrient availability in a perennial fruit crop

**DOI:** 10.1101/2021.03.11.434774

**Authors:** Ke Chen, David Kleijn, Jeroen Scheper, Thijs P.M. Fijen

## Abstract

Managing ecosystem services may reduce the dependence of modern agriculture on external inputs and increase the sustainability of agricultural production. Insect pollinators and arbuscular mycorrhizal fungi (AMF) provide vital ecosystem services for crop production, but it has not been tested whether their effects on crop yield interact and how their effects are influenced by nutrient availability. Here we use potted raspberry (*Rubus idaeus* L.) plants in a full-factorial randomized block design to assess the interacting effects of insect pollination, AMF inoculation and four levels of fertilizer application. AMF inoculation increased the per-plant flower number by 33% and fruit number by 35%, independently from insect pollination and fertilizer application. Single berry weight furthermore increased more strongly with fertilizer application rates in AMF inoculated plants than in non-inoculated plants. As a consequence, AMF inoculation boosted raspberry yield by 43% compared to non-inoculated plants. AMF inoculation increased pollinator visitation rate under intermediate fertilizer levels, suggesting additional indirect effects of AMF on yield. Fruit yield of pollinated plants increased more strongly with fertilizer application rate than the yield of plants from which pollinators had been excluded. At maximum nutrient availability, the combined benefits of both ecosystem services resulted in a 135% higher yield than that of fertilizer-only treatments. Our results suggest that benefits of ecosystem services on yield can be additive or synergistic to the effects of conventional management practices. Intensive, high-input farming systems that do not consider the potential adverse effects of management on ecosystem service providing species may risk becoming limited by delivery of ecosystem services. Pro-actively managing ecosystem services, on the other hand, has the potential to increase crop yield at the same level of external inputs.

## 1. Introduction

Agriculture depends on a wide array of ecosystem services (Costanza *et al*. 1997; Klein *et al*. 2007), but agricultural inputs like fertilizer have adverse effects on the species providing those services and on the wider environment (Bakhshandeh *et al*. 2017). Ecological intensification has been put forward as a promising way to make agriculture more sustainable and reduce negative impacts on the environment (Bommarco *et al*. 2013; Kleijn *et al*. 2019). This approach proposes to manage for biodiversity to complement or (partially) replace external inputs with production-supporting ecosystem services. Although ecological intensification is increasingly being advocated by scientists and policymakers as an environmentally friendly way towards food security (Pywell *et al*. 2015; IPBES 2016), it is rarely adopted by farmers (Kleijn *et al*. 2019). Farmers manage complex agro-ecosystems, with the interplay of several agronomic and environmental factors shaping crop yield. Evidence that a single ecosystem service has a positive effect on crop yield may not be convincing enough for farmers to change their day-to-day practices (Dainese *et al*. 2019; Kleijn *et al*. 2019). Ecological intensification might be more appealing to farmers when multiple ecosystem services together can synergistically enhance crop yield. This requires insight in the effects of multiple ecosystem services on crop yield simultaneously, whether and how these services interact and how their benefits are influenced by conventional agricultural practices. However, we are only just starting to understand how multiple ecosystem services may interact (Garibaldi *et al*. 2018; Tamburini *et al*. 2019), and we know even less how these interactions are being influenced by agricultural management. Here we contribute to addressing this knowledge gap by examining the interacting effects of aboveground insect pollination and belowground arbuscular mycorrhizal fungi (AMF) inoculation on crop yield of raspberry (*Rubus idaeus* L.) and how this is affected by different fertilizer application levels.

AMF are able to form symbiotic associations with about 72% of all vascular terrestrial plants (Smith & Read 2010; Brundrett & Tedersoo 2018), including the majority of field crops (Plenchette *et al*. 2005). AMF provide a range of services to plants, such as facilitating mineral nutrient uptake (mainly phosphorus and nitrogen), enhancing disease resistance and stress tolerance, and improving soil structure (Smith & Read 2010; Chen *et al*. 2018). AMF colonization of crop plants can significantly increase crop yield (Zhang *et al*. 2019). However, current agricultural practices, such as high fertilizer inputs and tillage, are likely to inhibit AMF growth, and root colonization may currently be suboptimal in many agricultural systems (Jansa *et al*. 2006). Farmers may actively manage for increased AMF colonization through reduced tillage (Bowles *et al*. 2017), or by inoculating the soil or seedlings, but whether this is effective for crop yield is less studied (Tamburini *et al*. 2020). Interestingly, AMF may also have indirect effects on crop production as the presence of AMF in plant roots can moderate the behavior of other service-providing species groups. For example, Gange and Smith (2005) found that plants with AMF can significantly increase pollinator visit frequency, which indicates that AMF and pollinator service delivery may interactively shape crop yield (Wolfe *et al*. 2005; Saini *et al*. 2019). However, AMF may also provide disservices to the host plant’s growth and development, for example by reducing phosphor uptake (Smith *et al*. 2004). Whether the net balance of AMF inoculation is positive for raspberry crop yield, and how this varies under different levels of fertilizer application is unknown.

Pollinators are important ecosystem service-providers as they contribute to 35% of the global food production, and enhance yields in two-thirds of global crops (Klein *et al*. 2007). Pollination may alter a number of interrelated qualitative and quantitative yield parameters such as fruit/seed set and size (Bommarco *et al*. 2012; Fijen *et al*. 2018). However, the positive effect of pollination on a particular yield parameter does not automatically result in a higher total crop yield. For example, in sunflower (*Helianthus annuus* L.) increasing insect pollination can contribute to higher seed set but with smaller seeds (Tamburini *et al*. 2017) resulting in the same overall yield, probably because yield is constrained by other factors, such as nutrient availability (Garibaldi *et al*. 2018). Particularly for high-revenue fruit crops like raspberry (Daubeny & Kempler 2003), both yield quantity and quality are important for farmers. To make more reliable predictions of the benefits of ecological intensification for agriculture, it is therefore important to gain insight in how effects of insect pollination shape crop yield through these intercorrelated yield parameters, and how this is affected by other ecosystem services such as those provided by AMF, or management practices such as fertilizer application.

Here, we experimentally manipulated insect pollination, AMF inoculation and nutrient availability on raspberry crop plants in a full-factorial randomized block design to test the potential interactive effect on yield of AMF inoculation and insect pollination at different levels of fertilizer application which, to our knowledge, has not been studied before. The main objectives of this study were (i) to test the effects of AMF inoculation and fertilizer application rates on pollinator visitation, (ii) to examine the effects of pollination and AMF inoculation on five yield quality and quantity parameters and how their effects are influenced by fertilizer application, and (iii) to explore the pathways explaining the relationships among the variables. The insights obtained in our study may help advance our understanding of whether and how we can integrate different ecosystem service into farming practices to make agriculture more sustainable.

## 2. Materials and methods

### (a) Study system

We used raspberry as our study crop, which is an increasingly important fruit crop with a global production value of $1.5 billion in 2018 (FAO 2018). We used the cultivar ‘*Tulameen’*, which is among the most popular raspberry cultivars worldwide due to its high marketable quality, mainly the appearance and flavour (Aprea *et al*. 2009). It is a self-compatible cultivar, but high-quality fruit production nevertheless benefits from visitation by insect pollinators (Daubeny & Kempler 2003; Chen *et al*. 2021). The study was carried out on an experimental field of Wageningen University & Research in Wageningen, the Netherlands (51° 59’ 47” N, 5° 39’ 36” E; 780 mm mean annual precipitation, 9.4 °C mean annual temperature).

### (b) Experimental design

In August 2019, we purchased raspberry plants with a height of ca. 60 cm from a local fruit tree supplier. To ensure that all plants were exposed to the same soil conditions, we carefully washed away any soil adhering to the roots of raspberry plants prior to transplanting. Each plant was then planted into a 10-litre plastic pot (upper diameter 28 cm, holes in the bottom for drainage but covered with root cloth to minimize root growth out of the pot), and filled with un-sterilized former agricultural soil (SOM content: 1.95%, available N: 14.0 mg/kg, available P: 0.6 mg/kg, available K: 19.4 mg/kg). Soils were not sterilized to reflect real-world conditions in agricultural fields where plants can be colonized by AMF already present in the agricultural soil.

As our AMF treatment, we added either alive inoculum (inoculated) or sterilized inoculum (non-inoculated). We used the commercially available *Rhizophagus intraradices* inoculum (MYKOS^®^ Xtreme Gardening, Canada). To sterilize the inoculum for our non-inoculated treatment, we autoclaved it at 121 °C for two hours (Changey *et al*. 2019). During transplantation, we gave each plant two tablespoons of inoculum or sterilized inoculum spread evenly on the roots.

The fertilizer treatments comprised four levels: 0, 33, 66 and 99 kg ha^−1^ of N per year. The fertilizer levels were selected to include the range from no to optimum N inputs, as the recommended annual fertilizer N application rates range from 45 to 85 kg/ha (Strik 2005). The annual dose was divided into three applications: the first one-third two weeks after transplanting (October 30, 2019), the second one-third at bud break (March 16, 2020) and the last one-third just before flower opening (April 24, 2020). We selected a local commonly used fertilizer for the experiment, containing 10.80% N, 13.44% K, 5.89% P, and 7.20% S (CropSolutions Co., Perth, UK).

This site is known to host pollinators, mainly wild bumblebees and managed honey bees, in sufficient densities to result in an optimal fruit set of raspberry plants (Chen *et al*. 2021). To examine the effect of insect pollination, we excluded pollinators from half of the plants and used open-pollinated plants as positive controls. We covered every plant of the pollinator exclusion treatments with a white semi-transparent mesh bag (mesh size 0.1 mm) before the onset of flowering and kept plants covered throughout the flowering period. The mesh bags allowed wind pollination but excluded all insect visitors. To avoid predation of the developing fruits, we covered all plants after flowering with the mesh bags until harvest.

We used a complete randomized block design with AMF (two levels), pollination (two levels) and fertilizer (four levels) fully crossed to measure their individual and interacting effects on raspberry productivity. This resulted in 16 treatment combinations, which were randomly assigned to individual raspberry plants and replicated in five blocks, bringing the total to 80 experimental plants. Potted plants were spaced one meter apart both within and between rows and dug into the soil to protect the roots from extreme temperatures. All plants received equal and ample irrigation, and weeds were regularly removed by hand.

### (c) Measurements

For each plant of the open pollination treatment, we conducted ten-minute pollinator censuses from May 12 to 27^th^ to see if the AMF and fertilizer treatments affected the pollinator visitation rate. We randomly observed plants ten times on different days (morning or afternoon), and only during sunny or slightly cloudy days and with low wind velocity, following the focal point observation method (Fijen & Kleijn 2017). We only recorded flower visitors that contacted anthers or stigmas of flowers. All flower visitors were identified on the wing.

From June 15 onward, we harvested ripe berries every other day and weighed each berry. Additionally, we counted the wilted and aborted flowers of each plant.

### (d) Data analysis

Four plants died over winter prior to fruit production, resulting in a dataset for 76 plants (Supplementary Table 1). Prior to analyses, single berry weight was averaged per plant to avoid pseudoreplication. Total flower number per plant was calculated as the sum of the total fruit number and the total number of flowers that did not develop into fruits (e.g. wilted or aborted flowers). Per-plant fruit set was calculated by dividing the fruit number by the total flower number and expressed as a percentage.

We fitted linear mixed-effects models to quantify the relations between the experimental treatments and response variables. We fitted separate full models for each of the response variables flower number, fruit number, fruit set (%), single berry weight (g/fruit) and total yield (g/plant), and included “block” as a random factor in all models. Independent variables included pollination, AMF inoculation, fertilizer application rate and their interactions. We also included a quadratic term for fertilizer application rate to test for non-linear relations between fertilizer levels and raspberry production (Tamburini *et al*. 2017). The full models were simplified by removing non-significant predictors (backward elimination) using likelihood ratio tests (Zuur *et al*. 2009). We additionally tested the effects of AMF and fertilizer treatments on average flower-visitor visitation rate (visitors/10 minutes), including the quadratic term for fertilizer application rate, and their interactions, and “block” as a random factor. For this analysis we only used the open pollination treatment plants. The models were built using the function lme() in the nlme package with the maximum likelihood estimation method (Pinheiro *et al*. 2019). Statistical assumptions of normality and homoscedasticity of model residuals were inspected visually through diagnostic plots. All analyses were performed in R (R Core Team 2020).

## 3. Results

### (a) Total visits and flower visitation rate

Altogether, 682 individual pollinators were observed, divided over seven taxa: *Apis mellifera* (471 individuals), *Bombus terrestris* congl. (132 individuals, cf. Williams *et al*. (2012)), *B. pascuorum* (55 individuals), *B. lapidarius* (13 individuals), *B. pratorum* (7 individuals), hoverfly (3 individuals) and *B. sylvestris* (1 individual). AMF inoculation and fertilizer application interactively influenced pollinator visitation rate (Table 1). Flower visitation rate increased with fertilizer levels, and was higher for plants that had been inoculated with AMF than for non-inoculated plants at intermediate fertilizer application rates, but not at low or high fertilizer application rates (Table 1, Fig. 1). Besides, flower visitation rate was strongly correlated with the number of flowers per plant (Supplementary Fig. 2).

**Table 1.** Effects of arbuscular mycorrhizal fungi (AMF; inoculated vs non-inoculated), pollination (open-pollinated vs pollinators excluded) and fertilizer application rates (0, 33, 66, 99 kg N·ha^−1^·year^−1^) on flower visitation rate (open-pollinated plants only, n=37) and raspberry fruit production variables (n=76). All analyses were performed using linear mixed-effects models. Bold values represent significant effects (P<0.05).

|  | Flower visitation rate |  | Flower number |  | Fruit set |  | Fruit number |  | Single berry weight |  | Yield |  |
| --- | --- | --- | --- | --- | --- | --- | --- | --- | --- | --- | --- | --- |
| | $\chi^2_{(1)}$ | P | $\chi^2_{(1)}$ | P | $\chi^2_{(1)}$ | P | $\chi^2_{(1)}$ | P | $\chi^2_{(1)}$ | P | $\chi^2_{(1)}$ | P |
| AMF | 2.096 | 0.148 | 8.074 | **0.004** | 0.007 | 0.933 | 5.436 | **0.020** | 0.277 | 0.599 | 7.712 | **0.005** |
| Pollination |  |  | 0.022 | 0.881 | 9.093 | **0.003** | 6.916 | **0.009** | 2.083 | 0.149 | 10.165 | **0.001** |
| Fertilizer | 5.394 | **0.020** | 19.934 | **<0.001** | 0.944 | 0.331 | 14.059 | **<0.001** | 8.725 | **0.003** | 23.003 | **<0.001** |
| Fertilizer^2 | 0.396 | 0.529 | 1.807 | 0.179 | 2.277 | 0.131 | 2.600 | 0.107 | 0.885 | 0.347 | 1.186 | 0.276 |
| AMF:fertilizer | 0.284 | 0.594 | 0.290 | 0.590 | 0.309 | 0.578 | 0.002 | 0.966 | 4.146 | **0.042** | 1.170 | 0.279 |
| AMF:fertilizer^2 | 5.234 | **0.022** | 3.565 | 0.059 | 0.577 | 0.448 | 0.607 | 0.436 | 1.164 | 0.281 | 0.324 | 0.569 |
| AMF:pollination |  |  | 0.071 | 0.790 | 0.375 | 0.540 | 0.040 | 0.841 | 0.140 | 0.708 | 0.552 | 0.458 |
| Pollination:fertilizer |  |  | 0.054 | 0.817 | 8.517 | **0.004** | 4.699 | **0.030** | 0.390 | 0.532 | 8.705 | **0.003** |
| Pollination:fertilizer^2 |  |  | 0.686 | 0.407 | 0.616 | 0.432 | 0.116 | 0.734 | 0.790 | 0.374 | 0.229 | 0.632 |
| AMF:fertilizer:pollination |  |  | 0.350 | 0.554 | 3.412 | 0.065 | 0.577 | 0.447 | 0.025 | 0.874 | 0.231 | 0.631 |
| AMF:fertilizer^2:pollination |  |  | 0.174 | 0.677 | 1.218 | 0.270 | 0.026 | 0.873 | 3.228 | 0.072 | 0.339 | 0.560 |

**Fig 1.**
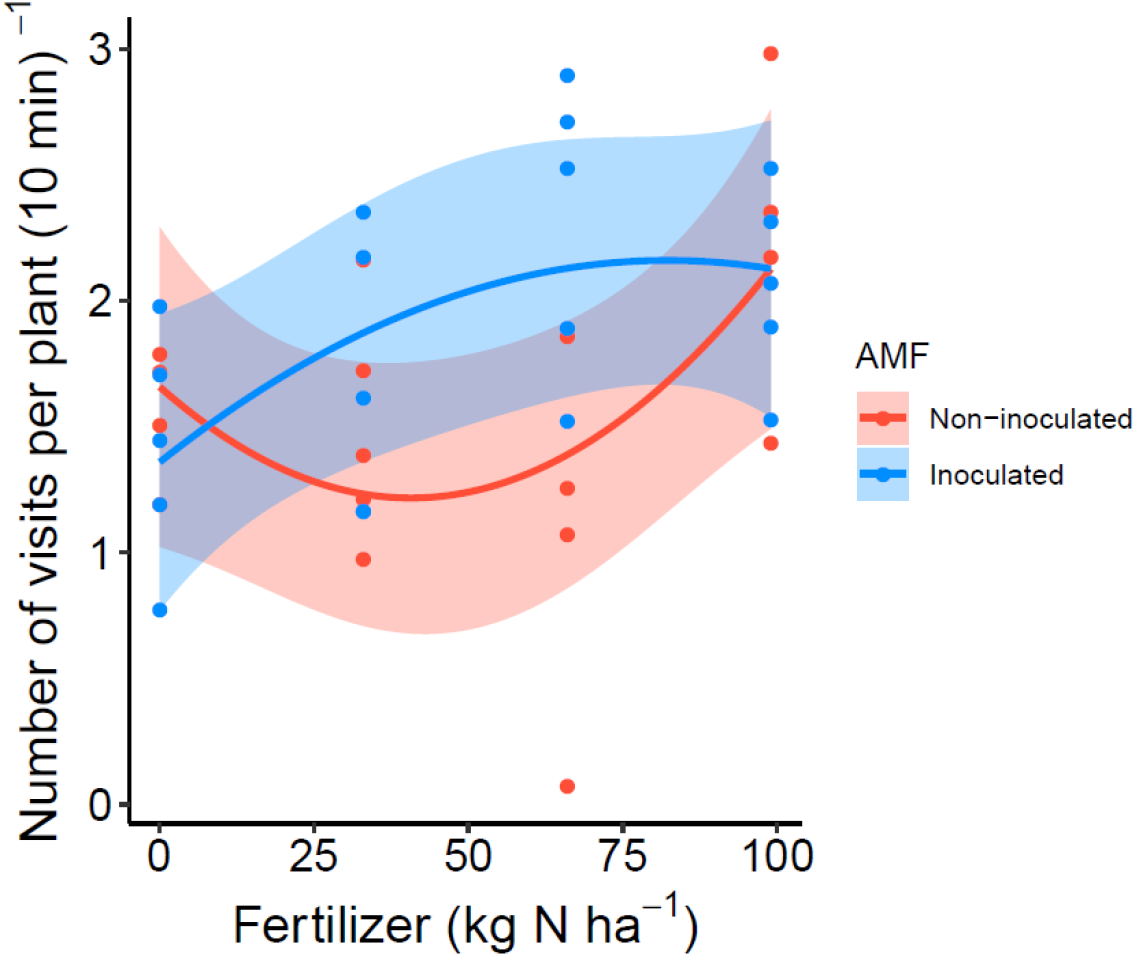
Interactive effects of AMF inoculation and fertilizer application rates on flower visitation rate (number of visits per 10 min) of raspberry. The lines are predicted by the minimum adequate model; shadings show the 95% confidence interval, and points represent partial residuals.

### (b) Flower number, fruit set and fruit number

The number of flowers per plant increased independently by both factors that (potentially) influence the nutrient acquisition, i.e. AMF inoculation and fertilizer inputs. Compared to the non-inoculated plants, AMF inoculation increased flower number by 33% (Fig. 2b, Table 1). Fertilizer inputs linearly increased flower number (Table 1), with plants receiving 99 kg N·ha^−1^ producing 105% more flowers than the unfertilized plants (Fig. 2a). There was a near-significant interaction (P=0.059) between the effect of AMF inoculation and the quadratic term of fertilizer application rate, with AMF inoculated plants receiving intermediate fertilizer application rates producing the most flowers (Table 1, Supplementary Fig. 1).

**Fig 2.**
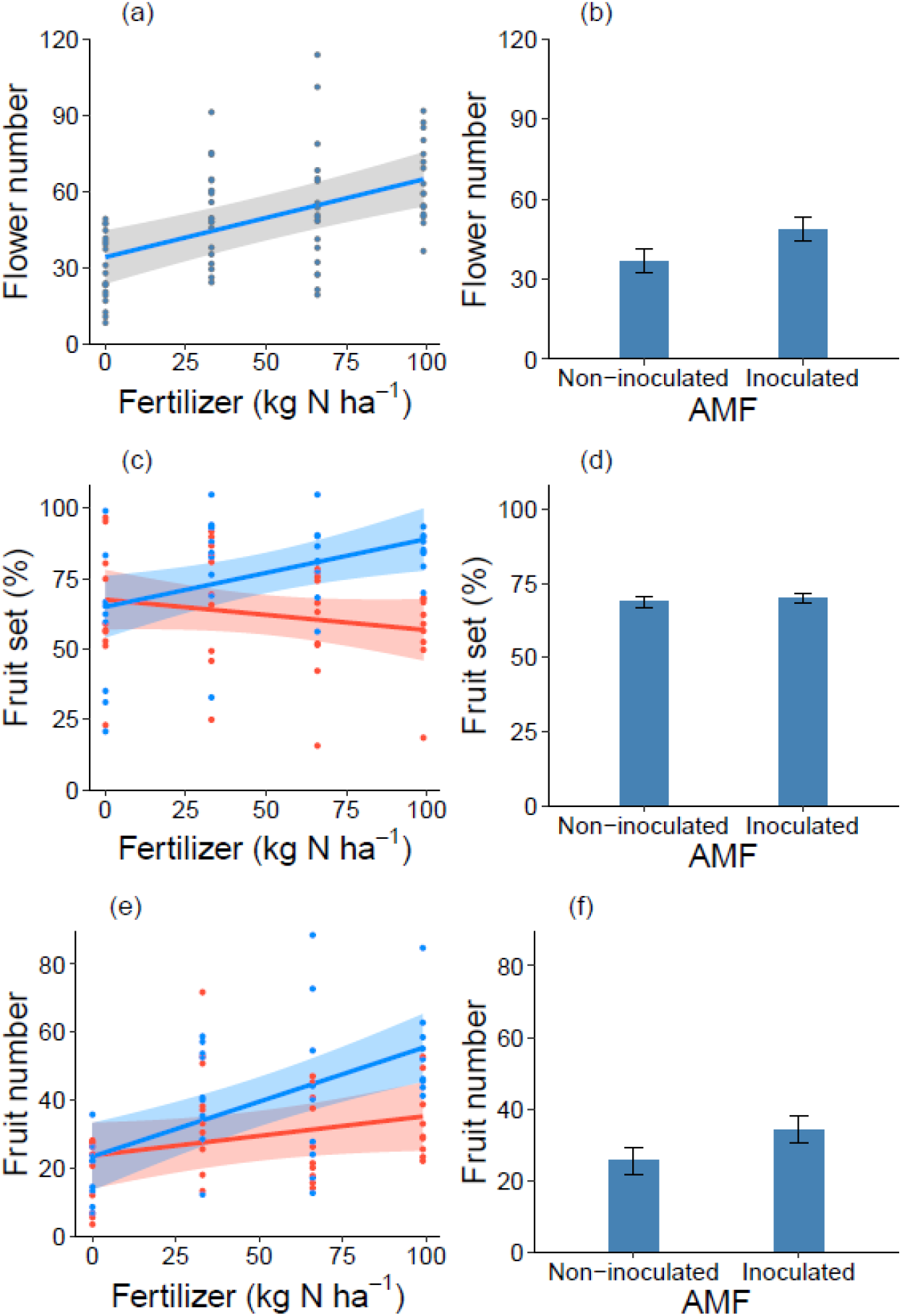
Effects of AMF inoculation, pollination and fertilizer application rates on flower number (a and b), fruit set (c and d), and fruit number (e and f) per plant. Pollination treatments are indicated by color in (c) and (e), pollinator excluded treatment in red and open pollination treatment in blue. Graphs show predicted values of the minimum adequate models; panel (d) shows non-significant estimated mean fruit set for AMF treatments as calculated in a model including AMF treatment (p=0.93) and the minimum adequate model parameters, and is shown for completeness. Shadings show the 95% confidence interval, and points represent partial residuals; error bars show ± 1 S.E.

Fruit set was mainly altered by insect pollination, but pollination benefits were most pronounced at the higher fertilizer application rates (significant pollination × fertilizer interaction; Table 1). From the lowest to the highest level, fertilizer application increased fruit set of open-pollinated plants by 37% and had little effect on fruit set in bagged plants (Fig. 2c).

Fruit number is the product of flower number and fruit set and this was clearly reflected in our results (Table 1; Fig. 2). AMF inoculation independently increased fruit number by 35% (Fig. 2f). Additionally, pollination and fertilizer application rate interactively affected fruit number with open-pollinated plants receiving 99 kg N·ha^−1^ producing 162% more fruits than unfertilized plants. This increase was only 53% when pollinators were excluded (Table 1; Fig. 2e).

### (c) Single berry weight and yield

Increasing fertilizer application rates influenced single berry weight interactively with AMF inoculation treatments, with a much more pronounced positive response in AMF inoculated plants compared to the non-inoculated plants (Table 1, Fig. 3). Pollination treatments did not significantly influence single berry weight (Table 1).

**Fig 3.**
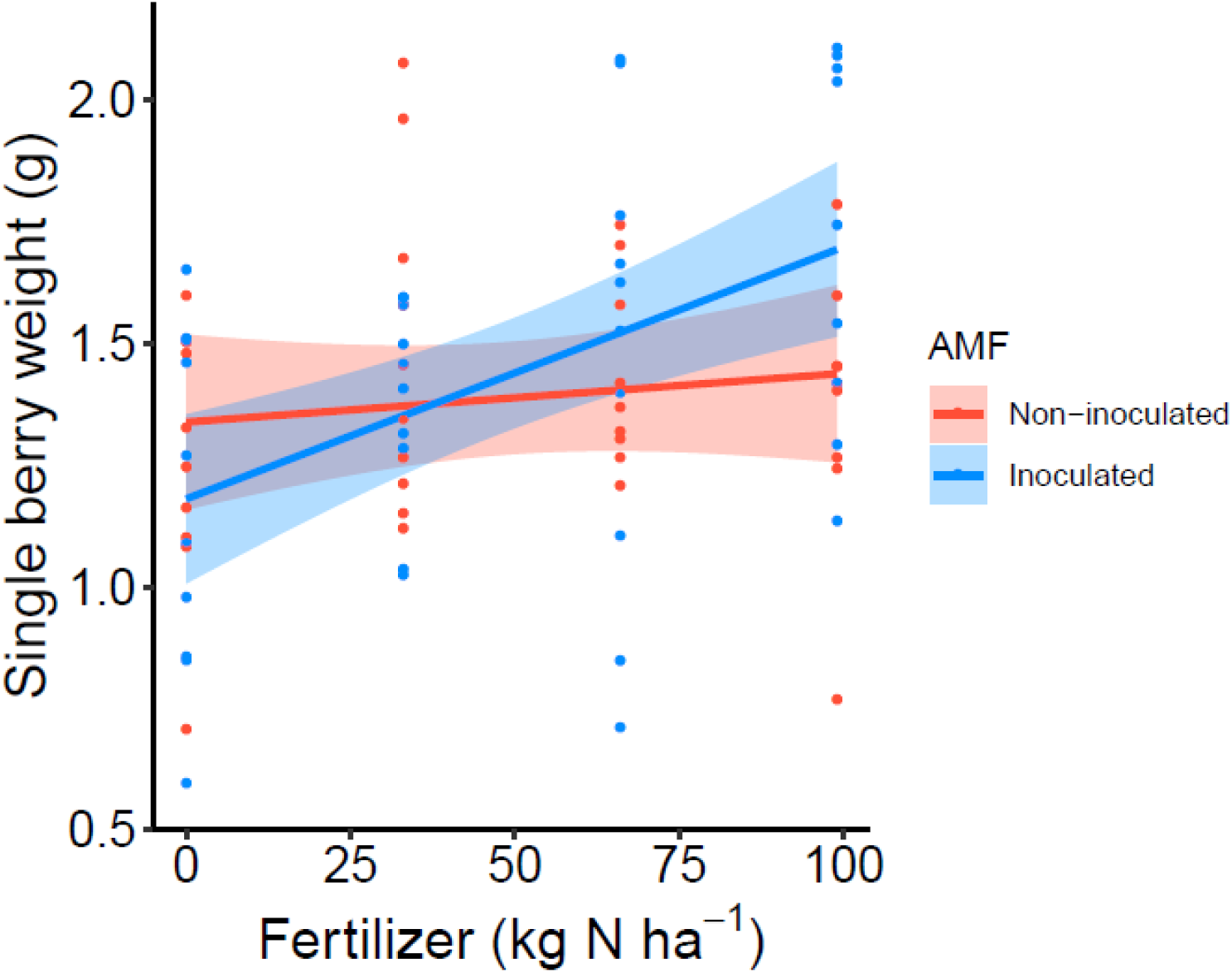
Interactive effects of AMF inoculation and fertilizer application rates on average single berry weight (g) per plant. The lines are predicted by the minimum adequate model; shadings show the 95% confidence interval, and points represent partial residuals.

The total yield is essentially the product of per-plant fruit number and single berry weight. However, total yield largely reflected effects of treatments on total fruit number, albeit stronger, while the significant interaction of AMF inoculation and fertilizer application on single berry weight was not reflected in the pattern for total yield (Table 1; Fig. 4). Total yield was positively related to fertilizer application rate, but these effects were much more pronounced in open-pollinated plants than in plants from which pollinators had been excluded; plants with insect pollination produced 90% more yield than bagged plants under our highest fertilizer input level. On top of that, the yield of AMF inoculated plants significantly increased by 43% compared to the non-inoculated plants (Fig. 4b). Under the highest fertilizer input, raspberry plants with open pollination and AMF inoculation produced the highest yield, on average 90.4 g berries, which was 135% more than the yield of plants receiving only the fertilizer application (38.5 g).

**Fig 4.**
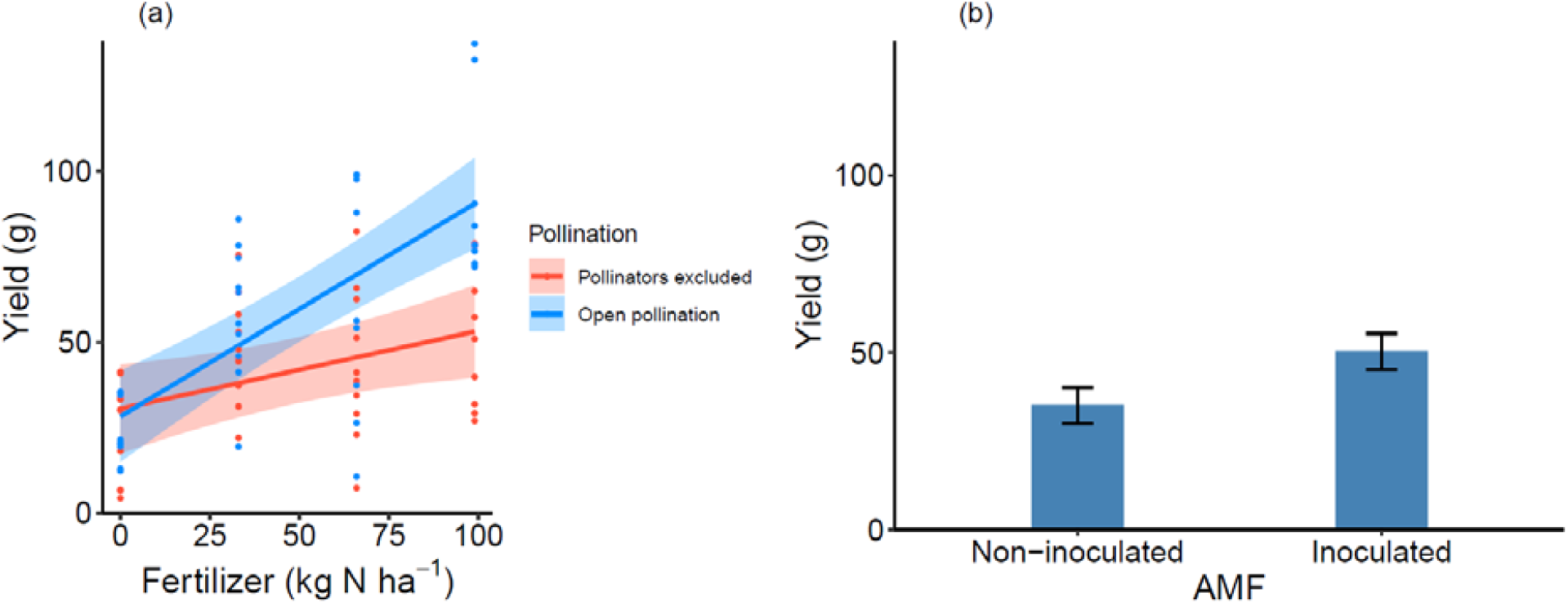
Effects of a) fertilizer application rates and pollination, b) AMF inoculation on yield per plant. Graphs show predicted values of the minimum adequate model (both); shadings show the 95% confidence interval, and points represent partial residuals (a); error bars show ± 1 S.E (b).

## 4. Discussion

Our results indicate positive effects of AMF inoculation on raspberry yield that were independent of the effects of pollination and fertilizer application, and positive synergistic effects of pollination and fertilizer inputs on yield. AMF inoculation enhanced the fruit-producing potential of plants by increasing the number of developed flowers on top of the already positive effects on the per-plant flower production of fertilizer. Pollination subsequently increased the likelihood that these flowers developed into fruits but only when plants received enough fertilizers. This probably suggests that poorly fertilized plants have insufficient resources for maximum fruit set. Interestingly, at intermediate fertilizer levels, AMF inoculation also enhanced pollinator visitation rates suggesting intricate indirect effects of one ecosystem service on another. Our findings imply that the simultaneous management of below- and aboveground ecosystem services can substantially increase the yield-enhancing effects of fertilizer application and represent a compelling example of ecological enhancement *sensu* Bommarco *et al*. (2013).

### (a) AMF inoculation contributing to raspberry yield directly and indirectly

AMF inoculation contributed to raspberry yield mainly through enhancing the number of flowers and by allowing plants to develop larger fruits. The 35% increase in fruit numbers of plants inoculated with AMF was very similar to the 33% increase in flower numbers of AMF inoculated plants, suggesting that AMF inoculation did not have a direct effect on fruit number but mostly on flower number. The effect on flower number may be due to the ability of AMF to increase plant nutrient concentrations (especially P and K) and to raise hormone levels stimulating bud-formation (Long *et al*. 2010) which have both been observed to lead to the development of larger numbers of flowers (Long *et al*. 2010). The positive effect of AMF inoculation on fruit size has been found in strawberry as well (Bona *et al*. 2015), but in our case the benefits were only expressed under ample fertilizer inputs (Fig. 3). Possibly, at low fertilizer application rates soil nutrient availability was the main limiting factor while at higher fertilizer application rates plant nutrient uptake capacity became a more limiting factor which AMF are known to improve. Surprisingly, when no fertilizer was applied, AMF-inoculated plants developed slightly smaller fruits than the plants that had not been inoculated, which could be the result of the competition for N with the host (Wang *et al*. 2018; Ingraffia *et al*. 2020). The interaction between AMF inoculation and fertilizer application did not carry over into final yield. Raspberry plants are readily colonized by AMF (Taylor & Harrier 2000) and it is to be expected that, regardless of treatment, all plants had formed associations with AMF to some degree by the end of the study. Our results therefore provide a conservative estimate of the potential contribution of AMF to raspberry crops.

Interestingly, our results indicate that AMF can also indirectly contribute to raspberry production through increasing pollinator flower visitation rate (Fig. 1) and thus pollination. Pollination has been shown to be an important factor limiting raspberry production, even in self-compatible cultivars like the one used in the present study (Chen *et al*. 2021). In our study, AMF and fertilizer inputs interactively shaped pollinator visitation rate (Fig. 1), and the pattern resembled their near-significant interaction on flower number (p = 0.059, Supplementary Fig. 1), which is an important plant trait to affect attractiveness to pollinators (Gange & Smith 2005). Therefore, it seems likely that the effects of AMF inoculation on pollinator visitation rate operated through their influence on flower number. However, we cannot rule out the possibility that AMF inoculation also influenced pollinator visitation rate through altering the composition of nectar and pollen (Somme *et al*. 2015; Bennett & Meek 2020).

### (b) Synergistic effects of insect pollination and fertilizer on raspberry production

Insect pollination and fertilizer inputs showed synergistic effects on raspberry yield and our results indicate that both are necessary for maximal yield (Fig. 4a). The possible pathway to explain the interacting effects starts with the positive effect of fertilizer on flower number, which simultaneously increased both the number of flowers that can potentially be pollinated and developed to fruits, as well as the attractiveness to pollinators (see Supplementary Fig. 2). Increased pollinator visitation rate generally enhances the transfer of pollen for ovule fertilization (Sáez *et al*. 2020), which may improve fruit set of the plants in the open pollination treatments (Fig. 2c). Interestingly, the benefits of insect pollination and fertilizer inputs seem to be depending on each other, as in the absence of the one, the benefits of the other diminish. For example, in the absence of fertilizer inputs, pollination benefits on fruit set are negligible, suggesting that nutrient availability limited the potential benefits of insect pollination to develop additional fruits (Garratt *et al*. 2018). Similarly, in the absence of insect pollination, solely increasing fertilizer inputs did not increase fruit set at all. This suggests that raspberry is probably limited by multiple ‘resources’ at the same time (Garibaldi *et al*. 2018), and that both need to be optimized to reach the highest raspberry crop yield. It also indicates that in our study system, ecosystem service benefits critically depend on the right management of external inputs and thus cannot easily replace them.

Because insect pollination did not influence single berry weight, the pollination-induced effects on fruit set carried over into similar effects on fruit number (Fig. 2e) and eventually yield (Fig. 4a). In a previous study using the same experimental system we did find positive effects of insect pollination on raspberry fruit size but not on fruit number (Chen *et al*. 2021). Plants have multiple ways to invest their most limiting resources (compensation mechanism; (Garratt *et al*. 2018)), which suggests that if one ecosystem service partially removes one limitation (e.g. nutrient-constrained flower development) this may impose new limitations to a subsequent process (e.g. nutrient-constrained drupelet development of raspberry fruits). However, it is noteworthy that regardless of the exact pathway, insect pollination resulted in substantially increased total raspberry crop yield in both studies.

### (c) The potential of capitalizing on ecosystem services in farming systems

Our results highlight the importance of maintaining ecosystem service providing species in agro-ecosystems. Not only did we find that without pollination and AMF inoculation raspberry yield would be substantially reduced, but yield effects of fertilizer were much less pronounced in the absence of ecosystem services. Agricultural production methods that do not consider potential adverse effects on ecosystem service providing species may risk shifting the system to one that is limited by delivery of ecosystem services rather than by management intensity (Deguines *et al*. 2014; Fijen *et al*. 2020). This is not a trivial issue as, for example, AMF colonization may be adversely affected by application of some types of pesticides (Hernández-Dorrego & Parés 2010; Hage-Ahmed *et al*. 2019). A farmer trying to control a disease using fungicides may succeed in minimizing disease damage only to lose the benefits provided by AMF. Our results furthermore suggest that pro-actively managing for ecosystem services can even increase crop production independently of conventional management practices such as fertilizer application, or can enhance the yield increases due to such practices as here with pollination. Such an approach could address the increasing demands for safe and healthy food that is typically associated with crop production methods that rely on natural processes rather than external inputs (Yiridoe *et al*. 2005). Here we found additive and synergistic benefits of both of the ecosystem service providing species groups that we examined. Given that other species groups can have additional yield impacts through, for example, biological pest control or nutrient cycling, the ultimate benefits to agricultural production of capitalizing more on natural processes could be substantially higher.

## Acknowledgements

We thank Lisa Kiel for her kind help with the pollinator observation. KC was funded by the China Scholarship Council (File No. 201706990023).

## Supplementary materials

**Supplementary Table 1.** The number of replicated raspberry plants survived in each treatment combination.

| Pollination | Fertilizer (kg·ha<sup>-1</sup><br>of N per year) | AMF | No. plants survived |
| --- | --- | --- | --- |
| Pollinators excluded | 0 | Non-inoculated | 5 |
| Pollinators excluded | 0 | Inoculated | 5 |
| Pollinators excluded | 33 | Non-inoculated | 5 |
| Pollinators excluded | 33 | Inoculated | 5 |
| Pollinators excluded | 66 | Non-inoculated | 5 |
| Pollinators excluded | 66 | Inoculated | 5 |
| Pollinators excluded | 99 | Non-inoculated | 5 |
| Pollinators excluded | 99 | Inoculated | 4 |
| Open pollination | 0 | Non-inoculated | 4 |
| Open pollination | 0 | Inoculated | 5 |
| Open pollination | 33 | Non-inoculated | 5 |
| Open pollination | 33 | Inoculated | 5 |
| Open pollination | 66 | Non-inoculated | 4 |
| Open pollination | 66 | Inoculated | 5 |
| Open pollination | 99 | Non-inoculated | 4 |
| Open pollination | 99 | Inoculated | 5 |

**Supplementary Fig 1.**
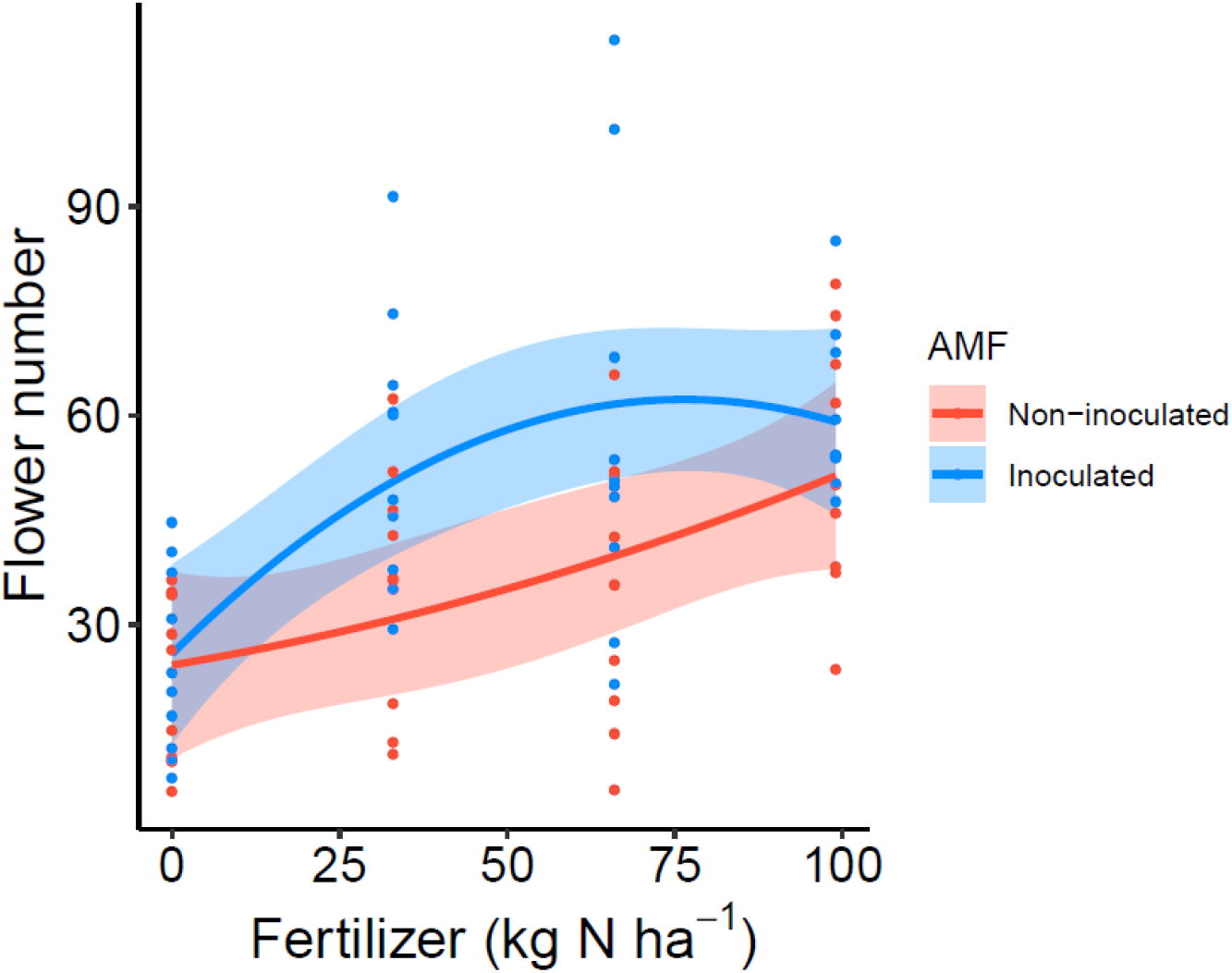
Interactive effects of AMF inoculation and fertilizer application rates on flower number per plant (near significant interaction, p=0.059). The lines are predicted by the minimum adequate model; shadings show the 95% confidence interval, and points represent partial residuals.

**Supplementary Fig 2.**
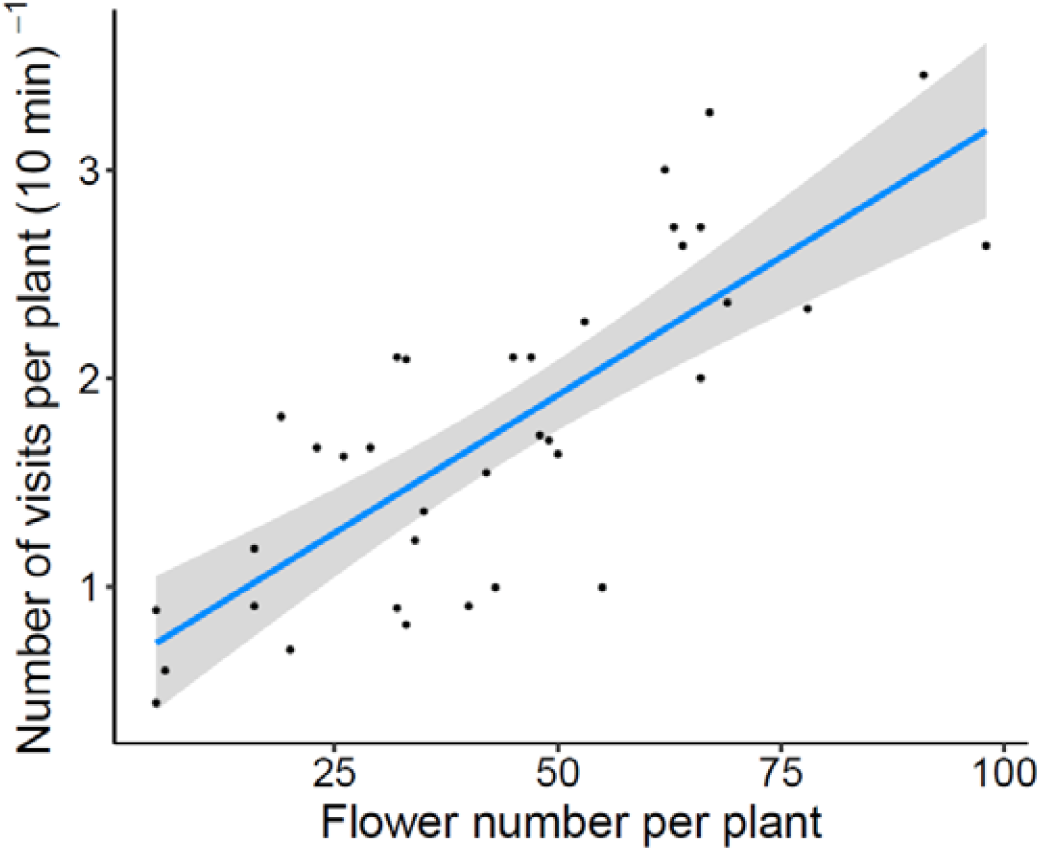
The relation between flower number and flower visitation rate (number of visits per 10 min) per plant, with the shading showing the 95% confidence interval. The graph bases on a simple linear regression model and the equation is y = 0.60 + 0.03x: (r^2^ = 0.61, p < 0.001)

## Reference

1. Aprea, E., Biasioli, F., Carlin, S., Endrizzi, I. & Gasperi, F. (2009). Investigation of Volatile Compounds in Two Raspberry Cultivars by Two Headspace Techniques: Solid-Phase Microextraction/Gas Chromatography− Mass Spectrometry (SPME/GC− MS) and Proton-Transfer Reaction− Mass Spectrometry (PTR− MS). Journal of agricultural and food chemistry, 57, 4011–4018.

2. Bakhshandeh, S., Corneo, P.E., Mariotte, P., Kertesz, M.A. & Dijkstra, F.A. (2017). Effect of crop rotation on mycorrhizal colonization and wheat yield under different fertilizer treatments. Agriculture, ecosystems & environment, 247, 130–136.

3. Bennett, A.E. & Meek, H.C. (2020). The influence of arbuscular mycorrhizal fungi on plant reproduction. Journal of Chemical Ecology, 46, 707–721.

4. Bommarco, R., Kleijn, D. & Potts, S.G. (2013). Ecological intensification: harnessing ecosystem services for food security. Trends Ecol Evol, 28, 230–238.

5. Bommarco, R., Marini, L. & Vaissière, B.E. (2012). Insect pollination enhances seed yield, quality, and market value in oilseed rape. Oecologia, 169, 1025–1032.

6. Bona, E., Lingua, G., Manassero, P., Cantamessa, S., Marsano, F., Todeschini, V. et al. (2015). AM fungi and PGP pseudomonads increase flowering, fruit production, and vitamin content in strawberry grown at low nitrogen and phosphorus levels. Mycorrhiza, 25, 181–193.

7. Bowles, T.M., Jackson, L.E., Loeher, M. & Cavagnaro, T.R. (2017). Ecological intensification and arbuscular mycorrhizas: a metalJanalysis of tillage and cover crop effects. Journal of Applied Ecology, 54, 1785–1793.

8. Brundrett, M.C. & Tedersoo, L. (2018). Evolutionary history of mycorrhizal symbioses and global host plant diversity. New Phytologist, 220, 1108–1115.

9. Changey, F., Meglouli, H., Fontaine, J., Magnin-Robert, M., Tisserant, B., Lerch, T.Z. et al. (2019). Initial microbial status modulates mycorrhizal inoculation effect on rhizosphere microbial communities. Mycorrhiza, 29, 475–487.

10. Chen, K., Fijen, T.P., Kleijn, D. & Scheper, J. (2021). Insect pollination and soil organic matter improve raspberry production independently of the effects of fertilizers. Agriculture, Ecosystems & Environment, 309, 107270.

11. Chen, M., Arato, M., Borghi, L., Nouri, E. & Reinhardt, D. (2018). Beneficial services of arbuscular mycorrhizal fungi–from ecology to application. Frontiers in Plant Science, 9, 1270.

12. Costanza, R., d’Arge, R., De Groot, R., Farber, S., Grasso, M., Hannon, B. et al. (1997). The value of the world’s ecosystem services and natural capital. Nature, 387, 253–260.

13. Dainese, M., Martin, E.A., Aizen, M.A., Albrecht, M., Bartomeus, I., Bommarco, R. et al. (2019). A global synthesis reveals biodiversity-mediated benefits for crop production. Science advances, 5, eaax0121.

14. Daubeny, H. & Kempler, C. (2003). ‘Tulameen’red raspberry. Journal of the American Pomological Society, 57, 42.

15. Deguines, N., Jono, C., Baude, M., Henry, M., Julliard, R. & Fontaine, C. (2014). LargelJscale tradelJoff between agricultural intensification and crop pollination services. Frontiers in Ecology and the Environment, 12, 212–217.

16. FAO (2018). Value of Agricultural Production. http://www.fao.org/faostat/en/#data/QV.

17. Fijen, T.P. & Kleijn, D. (2017). How to efficiently obtain accurate estimates of flower visitation rates by pollinators. Basic and Applied Ecology, 19, 11–18.

18. Fijen, T.P., Scheper, J.A., Boom, T.M., Janssen, N., Raemakers, I. & Kleijn, D. (2018). Insect pollination is at least as important for marketable crop yield as plant quality in a seed crop. Ecology letters, 21, 1704–1713.

19. Fijen, T.P., Scheper, J.A., Vogel, C., van Ruijven, J. & Kleijn, D. (2020). Insect pollination is the weakest link in the production of a hybrid seed crop. Agriculture, Ecosystems & Environment, 290, 106743.

20. Gange, A.C. & Smith, A.K. (2005). Arbuscular mycorrhizal fungi influence visitation rates of pollinating insects. Ecological Entomology, 30, 600–606.

21. Garibaldi, L.A., Andersson, G.K., Requier, F., Fijen, T.P., Hipólito, J., Kleijn, D. et al. (2018). Complementarity and synergisms among ecosystem services supporting crop yield. Global food security, 17, 38–47.

22. Garratt, M.P., Bishop, J., Degani, E., Potts, S.G., Shaw, R.F., Shi, A. et al. (2018). Insect pollination as an agronomic input: strategies for oilseed rape production. Journal of Applied Ecology, 55, 2834–2842.

23. Hage-Ahmed, K., Rosner, K. & Steinkellner, S. (2019). Arbuscular mycorrhizal fungi and their response to pesticides. Pest management science, 75, 583–590.

24. Hernández-Dorrego, A. & Parés, J.M. (2010). Evaluation of some fungicides on mycorrhizal symbiosis between two Glomus species from commercial inocula and Allium porrum L. seedlings. Spanish Journal of Agricultural Research, 43–50.

25. Ingraffia, R., Amato, G., Sosa-Hernández, M.A., Frenda, A.S., Rillig, M.C. & Giambalvo, D. (2020). Nitrogen Type and Availability Drive Mycorrhizal Effects on Wheat Performance, Nitrogen Uptake and Recovery, and Production Sustainability. Frontiers in plant science, 11, 760.

26. IPBES (2016). The assessment report of the Intergovernmental Science-Policy Platform on Biodiversity and Ecosystem Services on pollinators, pollination and food production. (eds. Potts, SG, Imperatriz-Fonseca, V & Ngo, HT). Secretariat of the Intergovernmental Science-Policy Platform on Biodiversity and Ecosystem Services, Bonn, Germany, p. 552.

27. Jansa, J., Wiemken, A. & Frossard, E. (2006). The effects of agricultural practices on arbuscular mycorrhizal fungi. Geological Society, London, Special Publications, 266, 89–115.

28. Kleijn, D., Bommarco, R., Fijen, T.P., Garibaldi, L.A., Potts, S.G. & van der Putten, W.H. (2019). Ecological intensification: bridging the gap between science and practice. Trends in ecology & evolution, 34, 154–166.

29. Klein, A.-M., Vaissiere, B.E., Cane, J.H., Steffan-Dewenter, I., Cunningham, S.A., Kremen, C. et al. (2007). Importance of pollinators in changing landscapes for world crops. Proceedings of the royal society B: biological sciences, 274, 303–313.

30. Long, L.-K., Yao, Q., Huang, Y.-H., Yang, R.-H., Guo, J. & Zhu, H.-H. (2010). Effects of arbuscular mycorrhizal fungi on zinnia and the different colonization between Gigaspora and Glomus. World Journal of Microbiology and Biotechnology, 26, 1527–1531.

31. Pinheiro, J., Bates, D., DebRoy, S., Sarkar, D. & Team, R.C. (2019). nlme: Linear and Nonlinear Mixed Effects Models.

32. Plenchette, C., Clermont-Dauphin, C., Meynard, J. & Fortin, J. (2005). Managing arbuscular mycorrhizal fungi in cropping systems. Canadian Journal of Plant Science, 85, 31–40.

33. Pywell, R.F., Heard, M.S., Woodcock, B.A., Hinsley, S., Ridding, L., Nowakowski, M. et al. (2015). Wildlife-friendly farming increases crop yield: evidence for ecological intensification. Proceedings of the Royal Society B: Biological Sciences, 282, 20151740.

34. R Core Team (2020). R: A Language and Environment for Statistical Computing. R Foundation for Statistical Computing.

35. Sáez, A., Aizen, M.A., Medici, S., Viel, M., Villalobos, E. & Negri, P. (2020). Bees increase crop yield in an alleged pollinator-independent almond variety. Scientific reports, 10, 1–7.

36. Saini, I., Aggarwal, A. & Kaushik, P. (2019). Inoculation with mycorrhizal fungi and other microbes to improve the morpho-physiological and floral traits of Gazania rigens (L.) gaertn. Agriculture, 9, 51.

37. Smith, S.E. & Read, D.J. (2010). Mycorrhizal symbiosis. Academic press. 38.

38. Smith, S.E., Smith, F.A. & Jakobsen, I. (2004). Functional diversity in arbuscular mycorrhizal (AM) symbioses: the contribution of the mycorrhizal P uptake pathway is not correlated with mycorrhizal responses in growth or total P uptake. New phytologist, 162, 511–524.

39. Somme, L., Vanderplanck, M., Michez, D., Lombaerde, I., Moerman, R., Wathelet, B. et al. (2015). Pollen and nectar quality drive the major and minor floral choices of bumble bees. Apidologie, 46, 92–106.

40. Strik, B.C. (2005). A review of nitrogen nutrition of Rubus. In: IX International Rubus and Ribes Symposium 777, pp. 403–410.

41. Tamburini, G., Bommarco, R., Kleijn, D., van der Putten, W.H. & Marini, L. (2019). Pollination contribution to crop yield is often context-dependent: A review of experimental evidence. Agriculture, ecosystems & environment, 280, 16–23.

42. Tamburini, G., Bommarco, R., Wanger, T.C., Kremen, C., van der Heijden, M.G., Liebman, M. et al. (2020). Agricultural diversification promotes multiple ecosystem services without compromising yield. Science advances, 6, eaba1715.

43. Tamburini, G., Lami, F. & Marini, L. (2017). Pollination benefits are maximized at intermediate nutrient levels. Proc Biol Sci, 284.

44. Taylor, J. & Harrier, L. (2000). A comparison of nine species of arbuscular mycorrhizal fungi on the development and nutrition of micropropagated Rubus idaeus L. cv. Glen Prosen (Red Raspberry). Plant and Soil, 225, 53–61.

45. Wang, X.-X., Wang, X., Sun, Y., Cheng, Y., Liu, S., Chen, X. et al. (2018). Arbuscular mycorrhizal fungi negatively affect nitrogen acquisition and grain yield of maize in a N deficient soil. Frontiers in microbiology, 9, 418.

46. Williams, P.H., Brown, M.J., Carolan, J.C., An, J., Goulson, D., Aytekin, A.M. et al. (2012). Unveiling cryptic species of the bumblebee subgenus Bombus s. str. worldwide with COI barcodes (Hymenoptera: Apidae). Systematics and Biodiversity, 10, 21–56.

47. Wolfe, B.E., Husband, B.C. & Klironomos, J.N. (2005). Effects of a belowground mutualism on an aboveground mutualism. Ecology Letters, 8, 218–223.

48. Yiridoe, E.K., Bonti-Ankomah, S. & Martin, R.C. (2005). Comparison of consumer perceptions and preference toward organic versus conventionally produced foods: A review and update of the literature. Renewable agriculture and food systems, 193–205.

49. Zhang, S., Lehmann, A., Zheng, W., You, Z. & Rillig, M.C. (2019). Arbuscular mycorrhizal fungi increase grain yields: A meta-analysis. New Phytologist, 222, 543–555.

50. Zuur, A., Ieno, E.N., Walker, N., Saveliev, A.A. & Smith, G.M. (2009). Mixed effects models and extensions in ecology with R. Springer Science & Business Media.

